# Environmental DNA reveals that rivers are conveyer belts of biodiversity information

**DOI:** 10.1101/020800

**Authors:** Kristy Deiner, Emanuel A. Fronhofer, Elvira Meächler, Jean-Claude Walser, Florian Altermatt

**Author notes:** Corresponding author K. Deiner.

## Abstract

DNA sampled from the environment (eDNA) is becoming a game changer for uncovering biodiversity patterns. By combining a conceptual model and empirical data, we test whether eDNA transported in river networks can be used as an integrative way to assess eukaryotic biodiversity for broad spatial scales and across the land-water interface. Using an eDNA metabarcode approach we detected 296 families of eukaryotes, spanning 19 phyla across the catchment of a river. We show for a subset of these families that eDNA samples overcome spatial autocorrelation biases associated with classical community assessments by integrating biodiversity information over space. Additionally, we demonstrate that many terrestrial species can be detected; thus revealing that eDNA in river-water also incorporates biodiversity information across terrestrial and aquatic biomes. Environmental DNA transported in river networks offers a novel and spatially integrated way to assess total biodiversity for whole landscapes and will transform biodiversity data acquisition in ecology.

“Eventually, all things merge into one, and a river runs through it.” — Norman Maclean

## Introduction

While rivers cover less than 1% of the landmasses on earth, they are invaluable for biodiversity and ecosystem services such as drinking water and energy production ^1^ Rivers, because of their characteristic dendritic network structure, also integrate information about the landscape through the collection and transport of sediments, organic matter, nutrients, chemicals and energy ^2,3^ For example, information contained in sediments allows us to understand how river drainages form and change in time as a result of climate and tectonic forces ^4^ Rivers also act as the lung of the landscape by releasing large fluxes of CO_2_ derived from terrestrial plant macromolecules, such as lignin and cellulose, through the breakdown and transport of coarse and fine particulate organic matter ^5^. River networks additionally play an important role in shaping patterns of genetic and species diversity for many organisms across the landscape by dictating dispersal pathways ^6,7^

Organic matter in the form of DNA is produced from organisms and is also transported through rivers via cells, tissues, gametes or organelles and is termed environmental DNA (eDNA) ^8–10^. DNA can be isolated from these organismal remains in the water, sequenced, and assigned back to the species of origin through the method of eDNA metabarcoding ^10,11^. This elegant process of collection and detection of a species’ DNA is becoming highly valuable for biodiversity sampling in ecology and conservation^10–17^. The spatial signal of eDNA, however, has only recently been explored and shows that in rivers eDNA can be transported over larger distances ^8,18^. Therefore, we hypothesized that rivers, through the aggregation and transport of eDNA, act as conveyer belts of biodiversity information which can be used to estimate species richness over broad spatial scales and across the land-water interface (Fig. 1a, Box 1).

**Figure 1.**
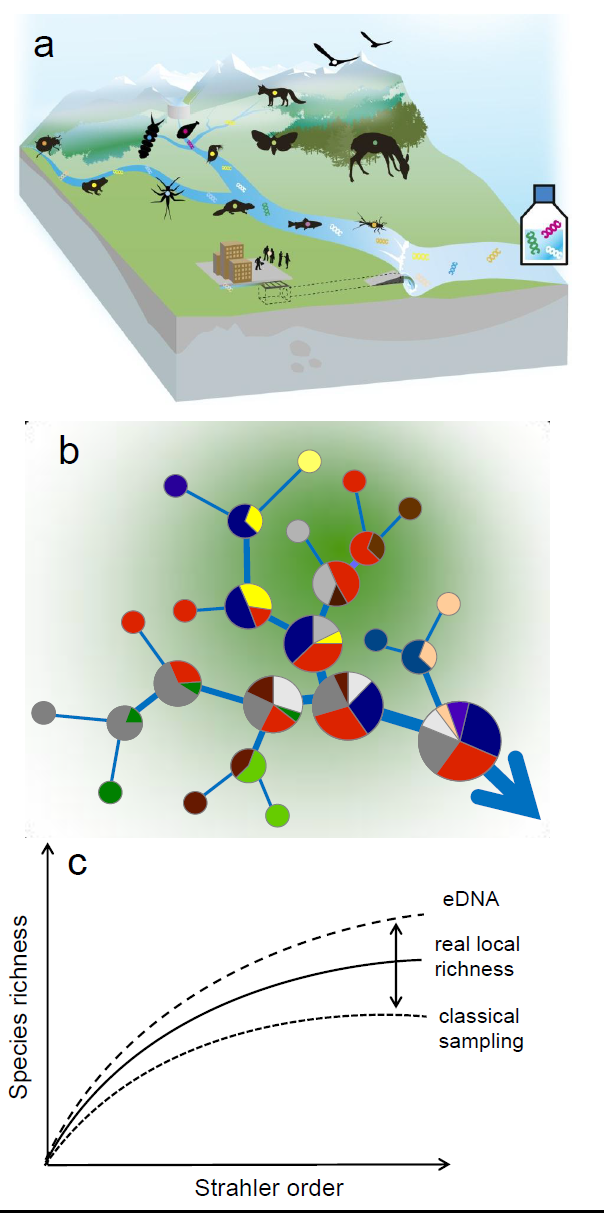
Conceptual model of environmental DNA spatial dynamics in a hypothetical river network. a) Visualization of species distribution in a landscape illustrating the release and accumulation of DNA in river water throughout its catchment. b) Characteristically high between-community diversity among headwaters (Strahler stream order 1; thinnest lines) is indicated by different colors representing local richness (α-diversity). Increasing size of pie chart indicates change in abundance. Flow direction is indicated with an arrow. Strahler stream order is indicated by the increasing width of river lines. c) While classical sampling only detects a fraction of real local diversity, eDNA sampling allows an estimate of catchment level diversity including both aquatic and terrestrial taxa and integrates this information across space due to downstream transport of eDNA.

The relevance of biodiversity sampling with eDNA found in river water is twofold. First, identifying biodiversity hotspots is invaluable for prioritizing global and regional conservation efforts ^19^. Estimates of richness to establish a place as a hotspot or not have suffered from being under-sampled ^20^. Under-sampling of biodiversity has many causes (and consequences) in conservation and ecology in general, but mainly comes from the sampling methods used for estimating richness in a way that is aggregated with respect to space ^21^. For example, a classical method for estimating richness of aquatic macroinvertebrates in rivers is to use a kicknet method, where all individuals in a certain defined area of a stream are collected in a net ^22^. Many such samples are then taken and subsequently pooled to represent richness for an entire river stretch or catchment. The pooling of spatially autocorrelated samples such as this causes an underestimation of biodiversity compared to if each species was independently sampled. Because it is typically infeasible to sample all species independently, statistical removal of the sampling artifact is recommended. Estimating biodiversity through eDNA is, however, a potential way to sample each species independent of space via their DNA becoming aggregated and transported through a river’s network (Fig. 1, Box 1).

Second, river biodiversity is highly affected by environmental changes and tracking these changes in space and time is of high interest ^23^. For example, the presence of tolerant (or absence of sensitive) aquatic organisms is important for determining water quality and has been used for over a century ^24–26^. This valuable metric known as ‘biomonitoring’ is entering a new era and the demand in its use has generated an undue burden on resource agencies. For example, the US, England, and Switzerland combined spend approximately 117.4 to 206.6 million US dollars annually on biomonitoring of aquatic systems (Table 1). This number represents only a small fraction of what countries spend on biomonitoring at more local levels, but characterizes the value we place on using species in their environment to monitor health of aquatic ecosystems.

**Table 1:** Annual bioassessment costs in millions of US dollars for freshwater resources. Information not available is abbreviated as NA. Original currency in brackets.

| Target Group | USA<sup>1</sup> | England<sup>2</sup> | Switzerland<sup>3</sup> |
| --- | --- | --- | --- |
| Fish | 31.4 - 58.2 | NA | 0.35 (0.33 CHF) |
| Benthic invertebrates | 38.1 - 70.7 | NA | 0.61 (0.58 CHF) |
| Algae (diatoms) | 34.7 - 64.5 | NA | 0.32 (0.3 CHF) |
| Macrophytes | NA | NA | 0.3 (0.29 CHF) |
| Total | 104.2 - 193.4 | 11.6 (7.3£) | 1.58 (1.5 CHF) |

Sources: <sup>1</sup> Stein *et al.* 2014; <sup>2</sup> Richard Walmsley, Forestry Commission personal communication;
<sup>3</sup> Markus Wüest, Federal Office for the Environment FOEN personal communication.

Biomonitoring is costly because of the different methods and expertise required to collect information about each targeted taxonomic group (e.g., Table 1) *^22,21^*. An eDNA method of biodiversity monitoring in rivers has several advantages in that it is non-lethal for most classically sampled taxonomic groups, minimizes habitat disruption, and can assess diversity across the tree of life with a single field sampling protocol making it extremely cost effective. Therefore, demonstrating the power of this tool to monitor biodiversity of important indicator groups (e.g., Table 1) in rivers will provide a fast, non-lethal and inexpensive alternative tool compared with classically used methods.

Whole community detection with environmental DNA has been called the ‘game changer’ for biodiversity sampling ^16^ and in this study we move this idea from theory into practice. We developed a conceptual model (Fig. 1; Box 1) and test the hypothesis that transported eDNA in rivers can be used in an unprecedented way to assess biodiversity of eukaryotes. We validate the ability of this method *in vitro* and *in situ* to assess globally important macroinvertebrate communities. We test whether taxonomic richness estimates from eDNA more accurately reflect the biodiversity of a rivers' catchment (Fig. 2), compared with classical methods, due to eDNA representing a spatially integrated signal. Lastly, we demonstrate that a large number of eukaryotic phyla from both aquatic and terrestrial taxa can be assessed from eDNA in river water and provide support for the hypothesis that rivers are conveyor belts of biodiversity information for landscapes.

**Figure 2.**
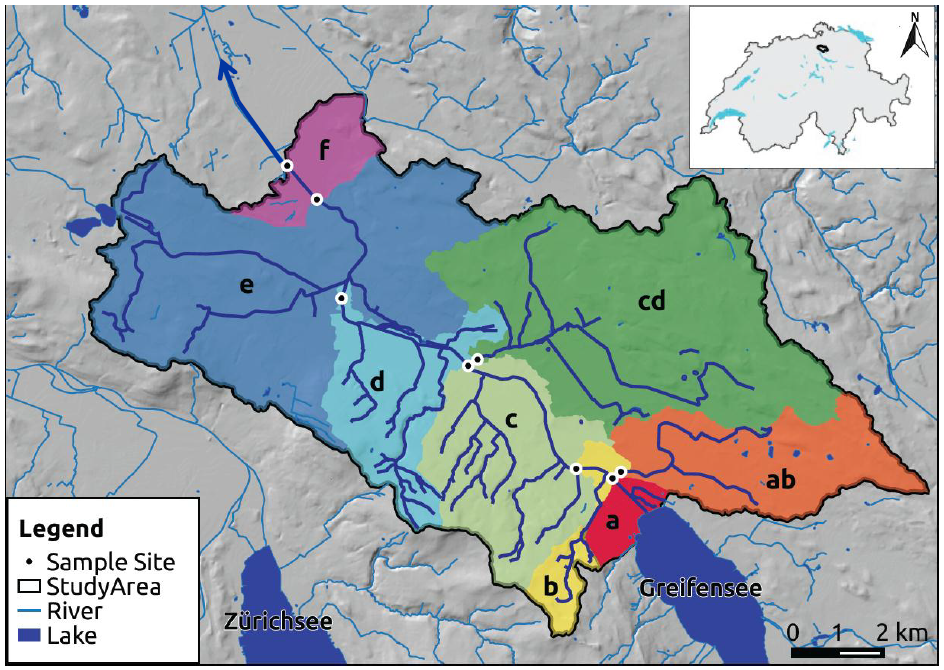
Study area and location of sampling sites where environmental DNA samples and classical sampling methods were carried out. The direction of flow for the river Glatt is northwest (blue arrow). The main stem of the river originates from the outflow of lake Greifensee. Colored regions represent the catchment upstream of each sampling point. Letters are used to indicate the position in the river network starting from the outflow ‘a’ to ‘f and the two sampled tributaries ‘ab’ and ‘cd’. Sources for GIS data were from Swisstopo (DHM25, Gewassernetz Vector 25) and reprinted with permission.

## Results

### All Eukaryotes

We detected a total of 296 families that span 19 eukaryotic phyla. All families were independently geographically verified as known to occur in Switzerland or the four neighboring countries (Fig. 3a, Supplemental Table 1). The majority of the families detected were Arthropoda (N = 196). Diversity in number of families detected was not proportional to read count and smaller organisms represented a much higher proportion of the sequences obtained (e.g., Rotifera; Fig. 3b). For example, two species in the phylum Rotifera accounted for 39% (92,907 sequences) of our data set. The majority of families were represented by more than ten sequences (N = 140, Supplemental Table 1). The largest data reduction step in the bioinformatic workflow was in linking a taxonomic name with our sequences (Supplemental Fig. 1, step E), resulting in only 4% (240,340 sequences) of acquired sequences that could be used for inferences in our study (Table 2). Of the sequences that were identified to species and that were independently geographically verified as occurring in Switzerland, many are terrestrial (N = 255, Fig. 4, Supplemental Table 2).

**Figure 3.**
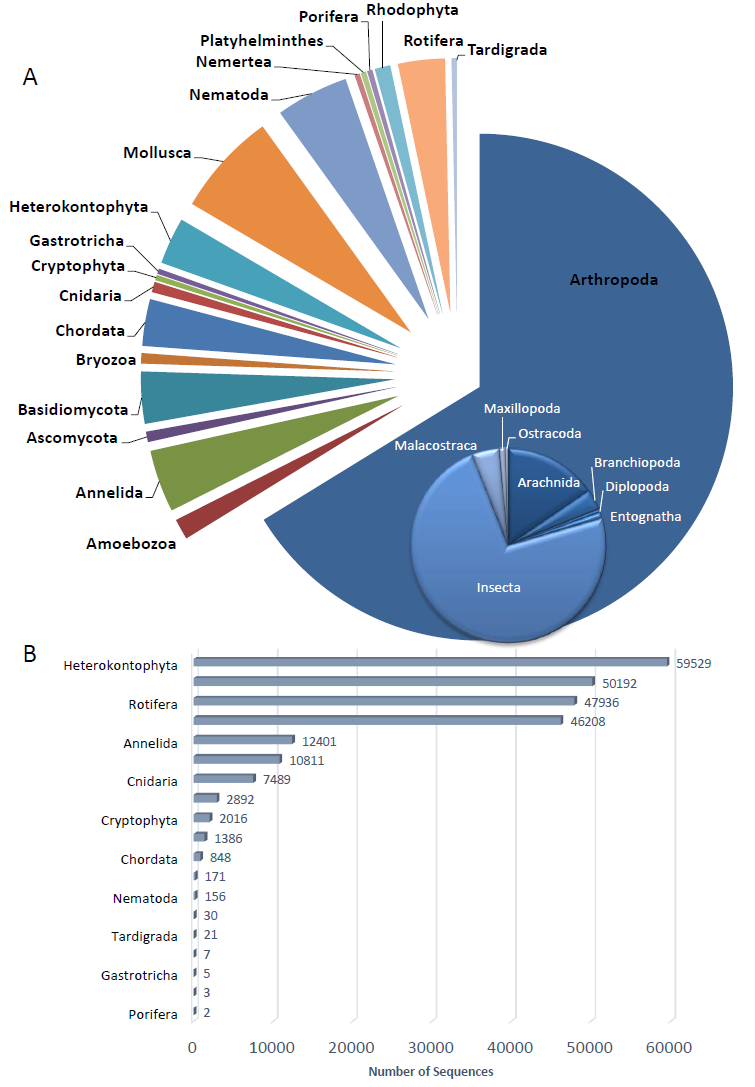
Total eukaryotic diversity detected from the river Glatt using environmental DNA metabarcoding. a) The number of families per phylum (N = 296) sampled and confirmed as known to be present in Switzerland or known from all four neighboring countries (Austria, France, Germany, and Italy). The inset further breaks the most abundant phylum (Arthropoda) into the number of families sampled by class (N = 196). b) Grey bars depict the number of sequences receiving a taxonomic identification at the family level for each phylum.

**Figure 4.**
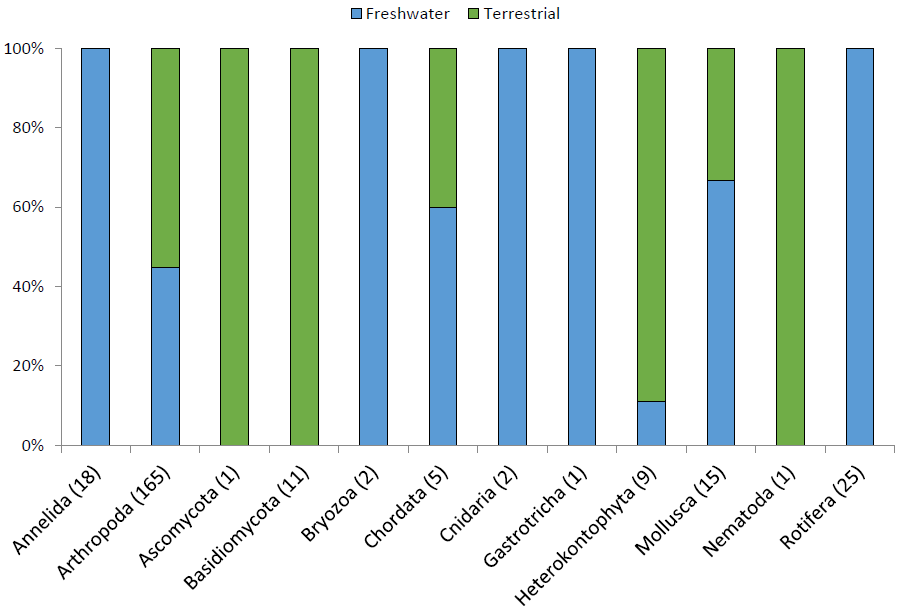
Percent terrestrial or freshwater species for the subset of each phylum that could be confirmed as known to be present in Switzerland or known from all four neighboring countries (Austria, France, Germany, and Italy; N = 255). Number in brackets indicates the number of species confirmed for each phylum.

**Table 2:** Sequences remaining after each bioinformatic filtering step and taxonomic assignment. Letter in parentheses of columns refer to bioinformatic filtering step with details given in methods and Supplemental Fig. 1. Assignment statistics are averages across all assignments for each site ± standard deviations (S.D.). Alignment length is in base pairs.

| Sample name | Site name | Raw read count | (A) Merged | (B) Quality Filter & Trim | (C) Mapping | (D) Chimera Removal | (E) Taxonomic Assignment | Average % ± S.D. of identical matches | Average ± S.D. reference alignment length |
| --- | --- | --- | --- | --- | --- | --- | --- | --- | --- |
| 02-01_S12 | a | 859156 | 496181 | 389859 | 251400 | 250725 | 32673 | 97.7 ± 2.7 | 204 ± 85 |
| 08-01_S18 | ab | 754781 | 479497 | 433506 | 151744 | 151422 | 21989 | 94.1 ± 2.7 | 195 ± 77 |
| 07-01_S17 | b | 587445 | 368095 | 304991 | 168274 | 167810 | 38316 | 98.1 ± 2.5 | 205 ± 81 |
| 03-01_S13 | c | 458635 | 294565 | 247769 | 144850 | 144453 | 37659 | 97.9 ± 2.6 | 213 ± 85 |
| 01-01_S11 | cd | 770055 | 512041 | 411234 | 176874 | 176231 | 36509 | 94.9 ± 2.8 | 195 ± 74 |
| 04-01_S14 | d | 664475 | 444870 | 392865 | 187002 | 186445 | 21783 | 96.6 ± 3.2 | 197 ± 77 |
| 05-01_S15 | e | 673495 | 364002 | 330412 | 172045 | 171389 | 20376 | 95.8 ± 3.4 | 192 ± 76 |
| 06-01_S16 | f | 1238917 | 740208 | 621210 | 311638 | 310375 | 31035 | 96.1 ± 3.3 | 193 ± 76 |
| Total for samples | | 6006959 | 3699459 | 3131846 | 1563827 | 1558850 | 240340 | 96.6 ± 3.2 | 200 ± 80 |
| Mock Community | | 421868 | 92934 | 82218 | 67983 | 67983 | 57641 | 94.0 ± 3.6 | 300.8 ± 38 |

### Macroinvertebrates

Of the 296 families detected with eDNA for eukaryotes, 65 are used in the Swiss biomonitoring program ^28^. Thirteen additional families were detected by kicknet samples only, totaling 78 macroinvertebrate families detected among our sampling sites of the river Glatt (Supplemental Fig. 2). From eDNA we recovered between 23 and 40 families at each site (Supplemental Fig. 2). With the classical kicknet method we sampled 17 to 24 families at each site (Supplemental Fig. 2). Of the total 78 families detected, 33 were detected by both methods, and often at the same location (Supplemental Fig. 2). Of the remaining 45 families, 32 were only detected with eDNA and 13 where only detected with the kicknet sample. Eleven of these 13 families were detected in the eDNA dataset, but did not meet bioinformatic thresholds used for filtering assignment values (e.g., where below a 90% sequence similarity or an alignment length less than 100 base pairs, Supplemental Table 3). The two undetected families (Potamanthidae and Aphelocheiridae) likely had insufficient sequence data on GenBank for identification (Supplemental Table 3). Of the 32 families only detected with eDNA, eight have been found in previous sampling events over the eighteen years of monitoring (Supplemental Table 4) and an additional two (Molannidae, Notonectidae) are known to occur in lake Greifensee, which feeds into the river Glatt, but are not known from the river Glatt (Supplemental Table 4).

Family richness (α-diversity) increased as a function of cumulative catchment area sampled for eDNA whereas this was not observed for kicknet samples (F_1,6_ = 5.45, p = 0.058, r^2^ = 0.95, eDNA; F_1,6_ = 0.0001, p = 0.99, r^2^ = 0.92, kicknet) (Fig. 5a). The slopes of the family-area relationship were different (slope_kicknet_ = 0.0006; slope_eDNA_ = 0.1077; F_1,12_ = 29.87, p = 0.0001), and the y-intercept was higher for eDNA compared with kicknet (F_1,13_ = 25.99, p = 0.0002) (Fig. 5a). β-diversity in the form of community dissimilarity did not increase as a function of distance for eDNA (r = 0.02, p = 0.44), whereas for kicknet sampling we observed an increase in dissimilarity (β-diversity) as a function of distance between sampling sites (r = 0.52, p = 0.005) (Fig. 5b).

**Fig. 1.**
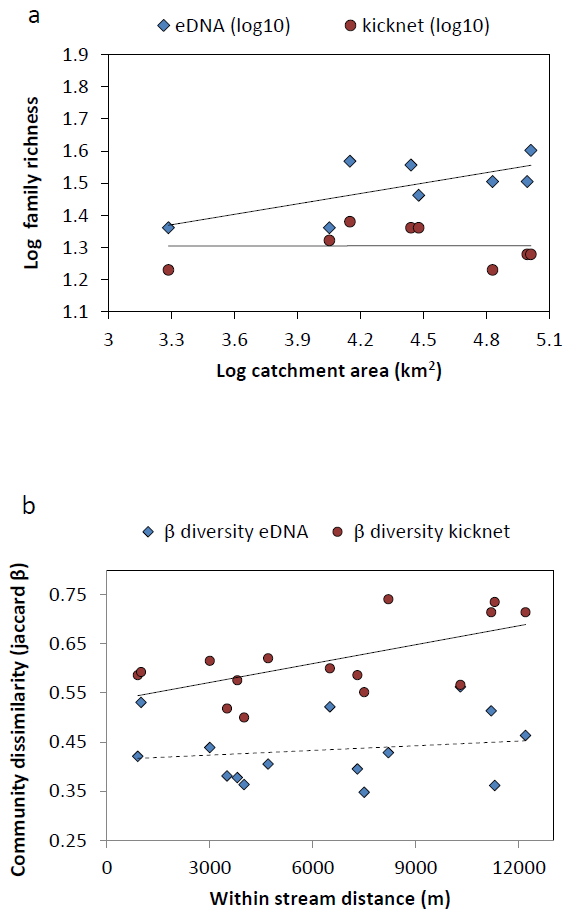
Difference of benthic macroinvertebrate family richness and community dissimilarity estimated between environmental DNA and kicknet sampling. a) α-diversity measured at each site and Log_10_ transformed taxon-area relationship for eDNA and kicknet samples. Slopes of line and the y-intercept are significantly higher for eDNA compared with kicknet (p<0.0001, p = 0.0002 respectively) indicating that eDNA samples a greater amount of diversity over a greater area compared with kicknets. b) Correlation of community dissimilarity with along-stream geographic distances between sample sites. Solid line for kicknet β-diversity indicates a significant positive relationship with stream distance (p = 0.005). Dashed line for eDNA β-diversity indicates no significant relationship with distance (p = 0.44).

### Mock communities

In total, we recovered 57,641 sequences from the mock community after the bioinformatic filtering and these sequences were identified to 25 of the 33 invertebrate taxa included in the mock community (Table 2, Supplemental Table 6). Of these sequences, 99.97% were correctly assigned to one of these taxa included in the mock community (Supplemental Table 7). The number of incorrectly assigned sequences was 0.03% (20/57,641) and all of these sequences belonged to two taxa (Tabanidae and Leuctridae, Supplemental Table 6). This resulted in a false positive rate of 8% (2/25). Increasing the stringency of our bioinformatics thresholds set for accepting an assignment to a level that removes all false positives (e.g., increasing assignment similarity to 92.1%) introduces false negative absences at a rate of 16% (4/25) in the mock community, that is, the exclusion of taxa that were present in the mock community but had assignment similarity less than 92.1 % (Supplemental Table 6).

## Discussion

We demonstrate that rivers, through their collection and transport of eDNA (Fig. 1), can be used to sample catchment-level biodiversity across the land-water interface. For aquatic macroinvertebrates, we found a greater richness in the number of families detected with eDNA compared with the classical kicknet method at the same sample location (Fig. 5a). This increased sensitivity is hypothesized to come from the process of transport of DNA through the network of a river. Transport of DNA through a river network decreases the biases associated with spatial autocorrelation (or limited scale of inference) inherent to classical kicknet community sampling (Fig.1; Box 1). The evidence from our work supports that eDNA found in rivers is a spatially integrated measure of biodiversity and this finding offers ecologists a new and unprecedented tool to sample landscape biodiversity with less sampling effort and potentially estimate richness of eukaryotic communities across biomes.

Our model (Box 1) identifies three important messages for the utility of eDNA as a genomic tool for biodiversity assessment. First, eDNA detection of species from river water decouples the presence of a species from its physical location in a habitat through downstream transport. Transport distance in empirical systems has been measured between 240 m up to 12 km ^8,29^, and thus allows for increased sensitivity in detection of patchily or elusively distributed species.

Additionally, transport of eDNA allows for richness estimates with less sampling effort because of the integrated signal over space. Second, eDNA will likely sample a higher diversity compared with classical sampling methods at any given site, but this depends on the local distribution of species and factors affecting transport and degradation of eDNA. Third, the interpretation of the species presence inferred from an eDNA sample in a river is different from that of classical sampling methods. Namely, eDNA detection of species should be interpreted as an integrated signal of presence and the spatial scale is determined based on the potential transport distance for a system. Thus, our model suggests that eDNA in rivers is an efficient tool for broad scale biodiversity assessments, and depending on the distance between water samples, less authoritative for very localized richness estimates.

Our data substantiate our conceptual predictions (Box 1), and by comparing eDNA with kicknet samples at each site, we highlight several important factors that illustrate both the power and current limitations of eDNA for biodiversity assessment. Many families of macroinvertebrates were detected at each site by both methods and have a great degree of overlap in which sites families were co-detected. For all sites, however, eDNA recovered more macroinvertebrate families compared to kicknet samples. We hypothesize this is likely due to the integrated signal from transported DNA which is evident by the fact that community composition does not change much (i.e., β-diversity remaining constant over distance), compared with kicknet estimated β-diversity which increased over the same river distance. This difference means that the two sampling methods give different information at the same site. Classical sampling methods give information that is localized, whereas the eDNA metabarcoding method in rivers measures presence of species on broader spatial scales. Scaling up of the classical community sampling method will likely always underestimate diversity ^21^, eDNA offers an empirical method to overcome this limitation and is an unparalleled way to estimate richness for larger areas. This novel finding is of great importance because in many cases estimating diversity for a large area is the goal, such as that for biodiversity hotspots^19^, conservation preserves ^20^, or entire river catchments ^30^.

Much of the current degradation of river habitat is at the catchment scale and cannot be attributed to a single point or source ^1^. Biomonitoring currently relies on the costly and lethal sampling of macroinvertebrates across many sites to understand ecosystem health of rivers (Table 1) ^27^ An eDNA signal of macroinvertebrates can be used to estimate more accurately diversity of a catchment with much less sampling effort. By contrast, understanding local changes in richness at a restoration site may still require classical sampling with kicknets. Interestingly, however, transport distances of eDNA are on a similar scale at which local species' pools are recognized to be important for recolonization of restored patches in a river system (0-5 km) ^31^. Therefore, eDNA could be used as a way to measure the species' pool available for recolonization. The scale of inference for eDNA, however, can be larger than 5 km due to long-distance transport within basins and between basins due to other vectors such as feces from predators. The complementarity between methods will aid in prioritizing river restoration efforts by identifying regions which have high recolonization potential of target species and possibly set expectations for the magnitude of change expected for restoration sites already in recovery.

Our results also identify a way of empirically measuring transport of community eDNA in rivers. Our analysis of β-diversity in this study system shows that community eDNA is likely transported and detected over a scale greater than 12 km. To determine the scale of transport for community eDNA in a river system, one subsequently needs to detect the scale at which there is a positive spatial autocorrelation between eDNA and β-diversity. This empirical measure of transport is needed because, as shown by our conceptual model, eDNA detection of biodiversity is a function of the transport distance, but also a function of the distribution of species within the network. Transport itself is furthermore affected by local factors, such as degradation of eDNA due to UV, pH and temperature ^32^, as well as discharge rates ^29^. Therefore, eDNA may not be necessarily transported and detected over the same distance for all river systems or consistently in time due to extreme events like heavy rainfall or drought. By using the correlation between β-diversity and river distance between sampling points, however, an *in situ* test can be performed and the scale of transport for community eDNA uncovered for any system and can be repeated across time to test if eDNA transport distance is stable in a system.

There are many important current limitations of the eDNA metabarcoding method. These challenges are related to factors such as the importance of primer or marker choice, the amplicon sequence length and the biodiversity detected as a function of the reference data available for identification of sequences ^33,34^ For example: fish, flatworms, and diatoms in our dataset are underrepresented to what we know occurs in the studied system. This is most likely due to the choice of primers, the genetic marker and the reference database. The primers used in this study are the universal Folmer primers for the 5’ end of COI ^35^, and it is known that these primers do not amplify DNA from fish and flatworms very well ^36,37; respectively^. Additionally, for diatoms it is known that COI is not the best genetic marker suitable for species level identification ^38^. Therefore, it is clear that more than one marker and/or primer set is needed to adequately assess biodiversity for the tree of life ^39^. However, use of an eDNA metabarcode method does not require additional sampling in the field. Rather it creates a single field sampling method whereby careful amplification of many genetic markers in the laboratory will enable an integrated detection for total biodiversity from a single sample.

An additional challenge faced by the further application of this approach is the need for continued development of diverse, but curated databases with taxonomically classified sequences. Our mock community analysis corroborated that we had a high accuracy in assignment of sequences when compared with the reference sequence generated from the DNA used for the mock community (98.4 to 99.9% similarity). This assignment accuracy dropped to (90.1 to 99.8%) when compared to NCBI's nucleotide database. From our eDNA samples we removed about 96% of the sequences and 15% of the loss was due to not being able to accurately assign a taxonomic name to the sequence. At the current filtering level applied here we are already accepting a false absence rate of 14% (Supplemental Table 3). Reducing our dataset further using more stringent criteria would increase type II error by creating many more false absences for taxa we actually collected in our kicknet samples at the time of sampling and thus know for sure they were present. Therefore, at this stage in deployment of an eDNA metabarcoding approach, we need to strive to reduce false absences balanced by knowing that false positive detections are also inevitable. In comparison with morphological assessments of macroinvertebrates at the family level, identification error is reported to range between 22.1% ^40^ and 33.8% ^41^, suggesting that the only alternative used in regulatory monitoring settings has a high false positive presence/absence rate. Most of the sequences from our dataset were removed because the taxonomic assignment failed. The solution for this is to increase deposition of sequences in curated databases ^42^ such as The Barcode of Life Database through continued collaboration between molecular ecologists and taxonomists. Digitizing specimens in the form of sequences is an essential step that will vastly improve our ability to accurately identify DNA found in the environment.

## Conclusions

We have demonstrated that rivers convey, through the collection and transport of environmental DNA, an unprecedented amount of information on biodiversity in landscapes. Our study shows that eDNA can be used to sample community structure of river catchments and do so even across the land-water interface. As such, detection of eukaryotic fauna with DNA found and transported in rivers may unite historically separated research fields of aquatic and terrestrial ecology and provide an integrated measure of total biodiversity for rapid assessment for one of the most highly impacted biomes of the world.

## Methods

### eDNA sampling, amplification and next generation sequencing

Water samples were collected from eight sites along the Glatt river network, a subcatchment of the Rhine River in Switzerland (Fig. 2). The study sites were chosen because they represent nodes in the river network where water from the major subcatchment tributaries combine and flow into the mainstem of the river Glatt. They also have a known history of monitoring macroinvertebrates for the past 15 years ^43^. At each site, DNA was isolated from between 840 to 900 mL of river water sampled. Method for sampling, capture and extraction of DNA followed that of Deiner *et al*. ^44^, where the capture method of filtration was coupled with a Phenol-Chloroform Isoamyl DNA extraction. Strict adherence to contamination control was followed using a controlled lab for eDNA isolation and pre-PCR preparations ^44^ Three independent extractions of 280 to 300 mL were carried out and then pooled to equal DNA captured and purified from 840 to 900 mL of water. Total volume of water filtered for each extraction replicate depended on the suspended solids in the sample of which clogged the filter. Water for this study was collected minutes prior to collecting aquatic macroinvertebrates using a classical sampling method kicknet, for description see below and ^43,45^ and therefore allowed for a comparison between the kicknet and eDNA methods for the detection of aquatic macroinvertebrate communities within the same watershed at the same time point.

Polymerase chain reactions (PCRs) were carried out for the target gene, cytochrome oxidase I (COI), using the universal COI primers ^35^ on pooled eDNA extractions for each of the eight sites and amplified a fragment of 658 base pairs (bp) excluding primer sequences. PCRs were carried out in 15 μL volumes with final concentrations of 1× supplied buffer (Faststart TAQ, Roche, Inc., Basel, Switzerland), 1000 ng/μL bovine serum albumin (BSA; New England Biolabs, Inc., Ipswich, MA, USA), 0.2 mMol dNTPs, 2.0 mMol MgCl_2_, 0.05 units per μL Taq DNA polymerase (Faststart TAQ, Roche, Inc., Basel, Switzerland), and 0.50 μMol of each forward and reverse primer ^35^. 2 μL of the pooled extracted eDNA was added. The thermal-cycling regime was 95 °C for 4 minutes, followed by 35 cycles of 95 °C for 30 seconds, 48 °C for 30 seconds and 72 °C for 1 minute. A final extension of 72 °C for 5 minutes was carried out and the PCR was cooled to 10 °C until removed and stored at −20 °C until confirmation of products occurred. PCR products were confirmed by gel electrophoresis on a 1.4% agarose gel stained with GelRed (Biotium Inc., Hayward, CA USA). Three PCR replicates were performed on each of the eight eDNA samples from our study sites and products from the three replicates were pooled. Negative filtration, extraction and PCR controls were used to monitor any contamination during the molecular workflow and were also replicated three times. Reactions were then cleaned using AMPure XP beads following recommended manufacturer's protocol except 0.6 × bead concentration was used instead of 1.8 × based on recommended protocol for fragment size retention of >500 base pairs (p. 31, Nextera XT DNA 96 kit, Illumina, Inc., San Diego, CA, USA). We quantified each pooled reaction using the Qubit (1.0) fluorometer following recommended protocols for the dsDNA High Sensitivity DNA Assay which has an accuracy for double stranded DNA between 0.005-0.5 pg/μL (Agilent Technologies, Santa Clara, CA, USA). At this step negative controls showed no quantifiable DNA and we therefore did not process them further.

The eight reactions were then each diluted with molecular grade water (Sigma-Aldrich, Co. LLC. St. Lewis, MO., USA) to 0.2 ng/μL following the recommended protocol for library construction (Nextera XT DNA 96 kit, Illumina, Inc., San Diego, CA, USA). Libraries for the eight sites were prepared using the Nextera XT DNA kit following the manufacturer's recommended protocols and dual indexed using the Nextera XT index kit A (Illumina, Inc., San Diego, CA, USA). Briefly, this protocol uses a process called tagmentation whereby the amplicon is cleaved preferentially from the 5’ and 3’ ends and the index and adaptor are ligated onto the amplicon. The tagmentation process produces an amplicon pool for each site (i.e., library) with randomly cleaved fragments averaging 300 bp in length that are subsequently duel indexed. The library constructed for each site was then pooled and paired-end sequenced (2 × 250 bp) on an Illumina MiSeq at the Genomic Diversity Center at the ETH, Zurich, Switzerland following the manufacturer's run protocols (Illumina, Inc., San Diego, CA, USA). The MiSeq Control Software Version 2.2 including MiSeq Reporter 2.2 was used for the primary analysis and the demultiplexing of the raw reads.

### Bioinformatic analysis

Workflow of process is presented in Supplemental Fig. 1. Run quality was assessed using FastQC version 0.10.1. Forward and reverse sequences were merged with a minimum overlap of 25 bp and minimum length of 100 bp using SeqPrep ^46^. Sequences that could not be merged were excluded from further analysis. Merged sequences with quality scores less than a mean of 25 where removed. Merged sequences were then de-replicated by removing exact duplicates, were de-noised using a sequence identity threshold of 99%, and were quality trimmed left and right by 28 bp using PrinSeq Lite version 0.20.3 to remove any primer sequence ^47^ Sequences were then mapped to the COI Barcode of Life Database (BOLD: iBOL phase 4.00) ^42^ using a map_reads_reference.py script with the minimum percent identity to consider a match as 50% and the minimum sequence length match to a reference of 50% to remove any sequences not likely of COI origin. Subsequent sequences were then chimera checked using usearch version 6 ^48^. Remaining sequences larger than 100 bp in length were then taxonomically identified using customized Blast searches against the NCBI non-redundant nucleotide database using the package blast 2.2.28, build on March 12, 2013 16:52:31 ^49^. Taxonomic assignment of a sequence was done using the best blast hit based on a bit score calculated using the default blastn search of a −3 penalty for a nucleotide mismatch and a reward of+1 for a nucleotide match. Sequences that did not match eukaryotes, were below 90.0% sequence similarity, had less than 100 bp overlap with query, had a taxonomic name not assigned below the level of family, matched best with unknown environmental samples and/or had a bit score less than 100 were excluded from biodiversity detection analysis for all sites. These parameters were used because they removed likely taxonomic identification errors or exclude data that was unidentified at the family level used for analysis ^44,50^. All raw sequences reads were deposited in NCBI's Sequence Read Archive (*accession numbers pending*).

After identification of sequences with the NCBI nucleotide sequence database, each uniquely identified taxon from any site was geographically verified as known to be present in Switzerland to the lowest level of taxonomy, or if no data was available for Switzerland, it was also considered present when the taxon was known to be present in Austria, France, Germany, and Italy. We excluded the one and very rare case (i.e., *Culicoides fascipennis)* where it is known for sure that a species is not in Switzerland, but found in all four neighboring countries. Geographic verification was done in consultation with 25 expert taxonomists for various groups, primary literature and through database repositories as described in Supplemental Table 1 and 2. If the species could be confidently confirmed as being present in Switzerland or in all four neighboring countries, their known habitat use was identified as being freshwater (defined as having at least one life stage inhabiting water), or terrestrial (which included species that inhabit riparian or wet habitats or typically feed in aquatic habitats, but do not have full life stages or reproduce in the water; Supplemental Table 2). Additionally, because we used BSA as an additive in PCR, we cannot rule out that detections of *Bos taurus* or *Bos indicus* were due to this reagent and therefore excluded them from analysis.

### Mock community analysis

A mock community approach was used to verify that our laboratory methods and bioinformatics pipeline were capable of correctly detecting the taxa of interest. We composed a mock community of invertebrate taxa from 33 different families spanning three phyla (all known to be present in our study area, Supplemental Table 5). We individually extracted their DNA, pooled and sequenced the mock community in accordance with the same methods used for analysis of eDNA samples from the river Glatt (see Supplemental Note for complete methods). We additionally Sanger sequenced all 33 DNA extractions from taxa following that of Mächler et al. ^51^ to generate a sequence reference database to assessing assignment errors when using NCBI's nucleotide database ^49^. All raw sequence files were submitted to NCBI's Sequence Read Archive (BioProject PRJNA291617) and details for each individual file are given in Supplemental Table 8.

### Kicknet sampling and identification

Macroinvertebrates were detected using a standard kicknet sampling design described for federal and cantonal guidelines in Switzerland ^28,45^ and represent our positive control for each site. Briefly, we took eight independent kicknet samples per site on October 29, 2012. Large inorganic and organic debris was removed and samples were pooled into a single collection jar with 70% EtOH. Jars were then stored at room temperature until morphological identification. This method and time of year has been shown to reflect the different microhabitats and provides a robust presence measure for many macroinvertebrates in Switzerland ^28^. Since eDNA has been shown to decay over short time periods of a few days to a few months; ^32^, using a single time point from a kicknet sample to compare with that of what is detected in the eDNA is valid. However, it is known that kicknet samples taken at different times of year, such as in the spring, can detect different species due to morphological constraints in the identification of specimens at young life stages or that their physical presence in the water is limited due to timing for their life cycle ^28^. Specimens from each site were sorted to the lowest taxonomic level possible (family, genus or species level) using dichotomous keys agreed upon by the Swiss Federal Office of the Environment ^28^. Specimens that could not be identified to at least to the taxonomic rank of family were excluded from further analysis.

### Comparison of eDNA and kicknet macroinvertebrate detection

For each site, we summarized the number of eDNA detected families of macroinvertebrates and number of families observed for the classical kicknet method including only aquatic taxa on the standardized list of macroinvertebrates for biomonitoring of Swiss waters by the Federal Office for the Environment ^28^. Using this standardized list, we calculated each site's observed α-diversity (local richness) for macroinvertebrates and visualized it on a heatmap of incidence. The estimated catchment area sampled for each position in the network was calculated as the cumulative sum of the area of all subcatchments into which all surface waters (excluding the lake) drain above the sampling point (Fig. 2). Topological distance between sampling sites was calculated along the river's path. Catchment area and distance between sampling sites was calculated using Quantum Geographic Information System in version 2.8 ^52^. The number of families detected (considered here as α-diversity) by each sampling method (eDNA and kicknet) was log_10_ transformed and regressed against the log_10_ of the river area to test for the taxon area relationship. We were interested in whether or not the two sampling methods differ in the magnitude of diversity detected due to the transport of DNA (y-intercept of the taxon area relationship), and that the rate of increase in number of taxa for a given area was faster for eDNA compared with the kicknet (slope of the regression lines) as predicted from our conceptual model. Slopes and y-intercepts of the two regressions for the taxon area relationship were tested using an analysis of covariance (ANCOVA).

To test for a spatial autocorrelation in community dissimilarity (*β*-diversity, using the Jaccard dissimilarity index) and between sampling locations we used a Mantel's test with 9999 permutations. Here we exclude the tributaries as it is not possible for eDNA to flow into these locations (e.g., cd into a). The Jaccard measure of *β*-diversity was used as it has been shown to estimate community dissimilarity for incidence data with less biases because of nestedness which is expected for the eDNA estimate of *β*-diversity due to transport ^53^. All statistical analyses were performed in R version 3.1.0 ^54^.

## Author contributions

K.D. & F.A. designed study, K.D., E.M. collected the data, J-C.W. developed the pipeline and conducted the bioinformatics analysis, E.F., F.A. and K.D. developed conceptual model, and all authors contributed to analysis and writing of the manuscript.

**Authors declare no competing interests**.

## Acknowledgements

We would like to thank Patrick Steinmann for assistance with kicknet samples and taxonomic identification, Katharina Kaelin for assistance with our study area map and geographic information used in this study, and Peter Penicka for contributing illustrations to Figure 1. We thank the 25 taxonomists who donated their time to help with the geographic verification of taxa identified through eDNA (all are listed in second worksheet of Supplemental Table 1 and 2). We thank Florian Leese, Vasco Elbrecht, Simon Creer and Michael Pfrender for providing insightful feedback on a previous version of the manuscript. Data analysed in this paper were generated in collaboration with the Genetic Diversity Centre (GDC), ETH Zurich. This work was funded by Eawag: Swiss Federal Institute of Aquatic Science and Technology and the Swiss National Science Foundation (grant no. PP00P3_150698 to FA).

## Box 1: Conceptual model exemplifying integration of biodiversity information along a river network using eDNA

Classical sampling methods, such as kicknet sampling, in rivers are very time- and cost-intensive (Table 1) ^27^Typically, sample methods for communities only capture a fraction of local a- diversity due to imperfect detection and sampling bias (Fig. 1c):

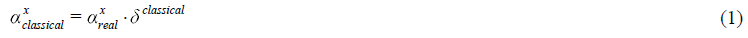

with 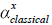 representing the measured α-diversity at a spatial location *x* in a river network using classical sampling methods, 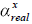 as the real α-diversity at this location and *δ*^classical^ as the detection rate of the sampling method. In order to comprehensively estimate the biodiversity of a river catchment, a large number of such samples is required. If samples are spatially autocorrelated, pooling of community samples will result in an underestimation of the real local richness ^21^.

Riverine networks, however, have the potential to collect this information for us ^2,3^ if we use an appropriate sampling method not biased by spatial autocorrelation for the area under study. Characteristic properties of rivers, such as the specific distribution of biodiversity ^45^ and transport of eDNA by the flow of water ^8^ make eDNA a promising method to estimate catchment level biodiversity while sampling at only one or very few locations

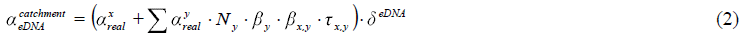

with 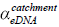 as the integrated measure of catchment α-diversity (see also Fig. 1). The sum captures the information integrated by the riverine system for all locations *y* (Strahler stream order) upstream of the sampling location *x.* The local diversity at a site of Strahler stream order *y* has to be weighted according to Horton's Law to capture the number of streams of this Strahler stream order (*N_y_*) ^3^ as well as by the Strahler stream order-characteristic *β*-diversity (*β*_*y*_). The estimate of catchment-level biodiversity increases with increasing β-diversity between the sampling point and all upstream locations (*β*_*xy*_) as well as with increasing transport distance (*τ*_*x*,*y*_; net rate including shedding and degradation). Note that the eDNA specific detection probability (*δ^eDNA^*) tends to be high as, in principle, only very few DNA molecules are needed for successful detection.

